# perfectphyloR: An R package for reconstructing perfect phylogenies

**DOI:** 10.1101/674523

**Authors:** Charith Bhagya Karunarathna, Jinko Graham

## Abstract

**Background:** A perfect phylogeny is a rooted binary tree that recursively partitions sequences. The nested partitions of a perfect phylogeny provide insight into the pattern of ancestry of genetic sequence data. For example, sequences may cluster together in a partition indicating that they arise from a common ancestral haplotype.

**Results:** We present an R package `perfectphyloR` to reconstruct the local perfect phylogenies underlying a sample of binary sequences. The package enables users to associate the reconstructed partitions with a user-defined partition. We describe and demonstrate the major functionality of the package.

**Conclusion:** The `perfectphyloR` package should be of use to researchers seeking insight into the ancestral structure of their sequence data. The reconstructed partitions have many applications, including the mapping of trait-influencing variants.

## Background

A perfect phylogeny is a rooted binary tree that represents a recursive partitioning of a set of objects such as deoxyribonucleic acid (DNA) sequences [1]. Though the perfect phylogenies are not ancestral trees, the structure of their nested partitions provides insight into the pattern of ancestry of DNA sequences. For example, the perfect phylogeny near a trait-influencing variant can provide useful information about trait association [2]. For instance, in a case-control study, case alleles may tend to cluster in a partition if the corresponding variant influences disease susceptibility. If a cluster has proportionally more case sequences than other clusters in the partition, there will be an association between the disease and cluster membership [3]. Thus, an R package to reconstruct perfect phylogenies from sequence data can be of use to researchers mapping the genetic location of trait-influencing variants.

We present an R package `perfectphyloR` to reconstruct perfect phylogenies underlying a sample of DNA sequences. The package uses a classic algorithm [1] together with heuristics [2] to partition sequences. Related software includes PerfectPhy [4] and BLOck aSSOCiation (BLOSSOC) [2].

PerfectPhy is a C++ program that implements efficient algorithms [5, 6] for reconstructing perfect phylogenies from multi-allelic DNA markers. The software comes with a collection of tools for importing/exporting files, handling missing data, filtering markers and drawing trees. PerfectPhy takes a given set of sequences and determines if it can be represented by a perfect phylogeny; if so, the partition is returned. The filtering tool can be applied in advance to select a maximal subset of markers compatible with a perfect phylogeny.

BLOSSOC is a C++ program for genetic fine-mapping that returns association statistics computed on perfect phylogenies. The statistics are calculated for moving windows of DNA markers across a genomic region of interest. The statistics are returned but not the partitions used to construct them. Unfortunately, BLOSSOC is no longer actively maintained (T. Mailund, personal communication) and is challenging to install on up-to-date operating systems.

Our package `perfectphyloR`, like BLOSSOC, is intended for use with moving windows of markers along the genome. The window sizes should be large enough to allow relatively fine partitioning of the sample of input sequences. However, requiring all the DNA markers in the window to be compatible with a perfect phylogeny tends to be too restrictive and leads to crude partitions. To avoid this limitation, we have incorporated the heuristics implemented in the partitioning algorithm of BLOSSOC. Since `perfectphyloR` returns the sequence partitions, users can then leverage any of the statistical and phylogenetic tools available in R to understand them. In addition, as an R package, the software is easier to install and to maintain as operating systems change.

Throughout, we assume the infinite-sites model and account for diallelic DNA markers only. Since our package reconstructs partitions regardless of whether the variants are common or rare, we refer to markers as single-nucleotide variants (SNVs) instead of single-nucleotide polymorphisms. By SNV, we mean any strictly diallelic marker. Our package is primarily directed to applications at the population level, rather than the interspecies level. Briefly, a neighborhood of SNVs is determined about a focal SNV, as described below. Then, the perfect phylogeny is built by recursive partitioning on SNVs in this neighborhood.

We first discuss the implementation of the reconstruction of the partitions underlying a sample of DNA sequences. We then illustrate the major functionality of the package with worked examples.

## Implementation

In this section, we describe the reconstruction process, which consists of three steps:

1. Create a `hapMat` data object.
2. Reconstruct the perfect phylogeny at a focal SNV.
3. Reconstruct perfect phylogenies across a genomic region.

We first create an object of (S3) class `hapMat` containing SNV sequences to be partitioned with the function `createHapMat()`. To construct a `hapMat` data object, users are required to specify:

- `hapmat`, a matrix of 0’s and 1’s, with rows representing sequences and columns representing SNVs,
- `snvNames`, a vector of names of SNVs labelling the columns of `hapmat`,
- `hapNames`, a vector of names labelling the sequences in the rows of `hapmat`,
- `posns`, a numeric vector specifying the physical locations along the chromosome (in base pairs) of SNVs in the columns of `hapmat`.

In principle, and as noted by a reviewer, the `hapMat` structure could be extended to accommodate multi-allelic variants, although we do not pursue this here.

With the main function `reconstructPP()`, the user can reconstruct the perfect phylogeny at a chosen focal SNV. The result is a `phylo` object to which the user may apply all the tools from the `ape` package [7] for summarizing the reconstructed partition of sequences.

The function `reconstructPP()` consists of three major steps:

1. Determine a neighborhood of SNVs around a given focal SNV.
2. Order the SNVs in the neighborhood.
3. Recursively partition sequences based on SNVs in the neighborhood.

For a given focal SNV, the algorithm finds a neighborhood of SNVs. Starting from the focal SNV, the neighborhood of SNVs that are compatible with the focal SNV is expanded as much as possible on either side of the focal SNV until an incompatible SNV is found. The compatibility of a pair of SNVs is determined by the Four-Gamete Test [8]. For example, under the infinite-sites mutation model and no recombination, if the patterns at two SNVs are 00, 01, 10 and 11, then a mutation must have occurred twice at the same SNV and the two SNVs are said to be incompatible. If the neighborhood of compatible SNVs is smaller than a user-defined minimum size, we include incompatible SNVs in order of their physical proximity to the focal SNV, until the minimum size is reached.

Once the neighborhood of SNVs is determined, we order the compatible SNVs in the neighborhood from the most ancient to the most recent based on the minor allele frequency. We use the minor allele frequency of an SNV as a proxy for its age. Our rationale is that, under the infinite-sites mutation model, the age of SNVs can be inferred from the derived allele frequency. Then, we order incompatible SNVs according to their physical proximity to the focal SNV.

The algorithm partitions sequences based on the most ancient compatible SNV in the neighborhood, and then recursively moves towards the most recent compatible SNV. When there are no further compatible SNVs in the neighborhood, the algorithm partitions sequences based on the incompatible SNVs, in order of their physical proximity to the focal SNV. Starting with the most ancient compatible SNV in the neighborhood, the algorithm partitions the sequences based on their carrier status for its derived allele. Then the algorithm jumps to the next-oldest compatible SNV in the neighborhood based on allele frequency and continues partitioning. After considering the compatible SNVs, the algorithm moves to any incompatible SNVs in the neighborhood in order of their physical proximity to the focal SNV. This process is repeated until each cluster contains only one sequence or there are no more SNVs to consider in the neighborhood. Thus, the method requires phased data. If a user has unphased data, phasing can be done in advance with software such as fastPHASE [9], BEAGLE [10], IMPUTE2 [11], or MACH [12, 13].

## Examples

This section gives worked examples illustrating how to reconstruct the partitions underlying a sample of DNA sequences. In addition, we show how to investigate the association between the reconstructed partitions and a user-specified partition. The association statistics we consider include the Rand index [14], the distance correlation (dCor) statistic [15], the Heller-Heller-Gorfin (HHG) statistic [16], the Mantel statistic [17], and the R-Vector (RV) coefficient [18]. The Rand index quantifies the association between two partitions directly. The dCor statistic, HHG statistic, Mantel statistic, and RV coefficient quantify the association between two distance matrices derived from partitions.

We first illustrate how to create a `hapMat` data object of SNV sequences. We then reconstruct a perfect phylogeny at a focal SNV. Next, we reconstruct perfect phylogenies across a genomic region. Finally, we show how to visualize and test associations between these reconstructed partitions and

- a comparator partition or dendrogram,
- a comparator distance matrix, and
- a phenotypic distance matrix.

To illustrate, we consider a toy example with 4 sequences comprised of 4 SNVs at positions 1, 2, 3, and 4 kilo-base pairs (kbp). The required `hapMat` object is created by executing the following command:

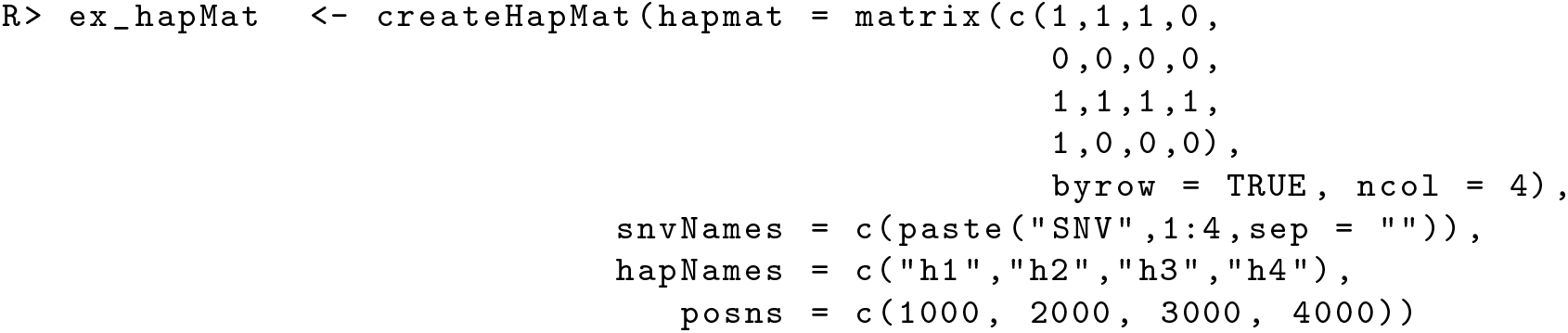

The structure of the resulting object of class `hapMat` is as follows.

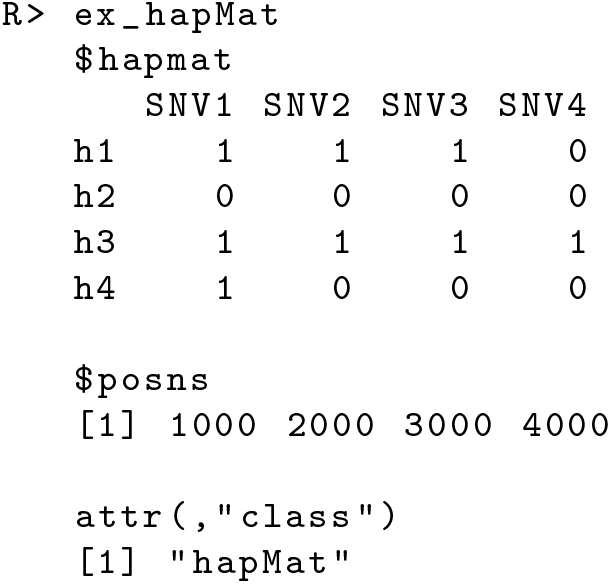

If a user has a variant call format (`vcf`) file that consists of SNV data with a single alternative allele and no missing values in the genotype field, the `hapMat` data object can be created by supplying the file path to the `vcf` file as follows:

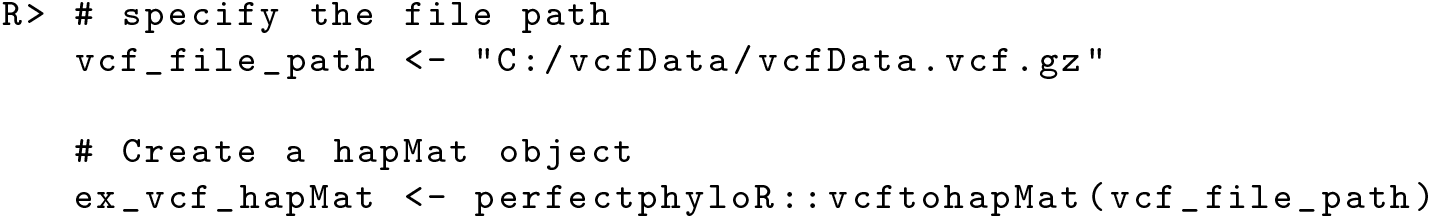

Once the `hapMat` object is created, the user can reconstruct a perfect phylogeny at a focal SNV with `reconstructPP()`, by specifying the following four arguments:

1. `hapMat`: A data structure of class `hapMat`, created by `createHapMat()`.
2. `focalSNV`: The column number of the focal SNV at which to reconstruct the perfect phylogeny.
3. `minWindow`: Minimum number of SNVs around the focal SNV in the neighborhood of SNVs used to reconstruct the perfect phylogeny (default is the maximum of one and 2% of the total number of the SNVs).
4. `sep`: Character string separator to separate sequence names for sequences that can not be distingiushed in the neighborhood around the focal point. For example, if sequences “h1” and “h3” can not be distinguished and `sep = “−”`, then they will be grouped together with the label “h1-h3”. The default value is `“−”`.

For example, consider the dataset `ex_hapMatSmall` data comprised of 10 sequences and 20 SNVs. This dataset is a subset of the larger example dataset, `ex_hapMat_data`, that comes with the package. The larger dataset has 200 sequences and 2747 SNVs, and was used in a previously published association association analysis [19]. We can reconstruct a perfect phylogeny at the first SNV of `ex_hapMatSmall_data` by executing the following commands:

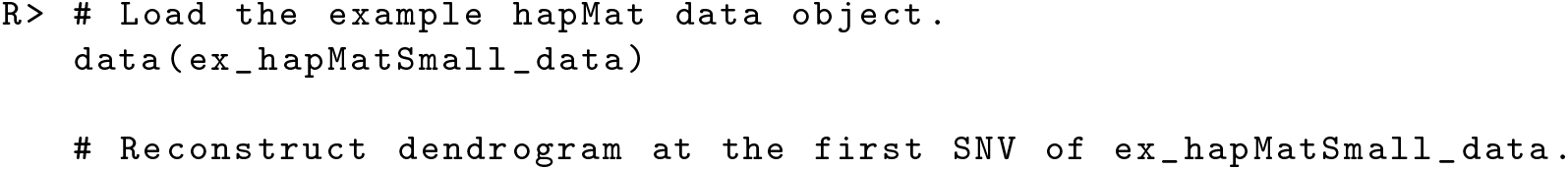

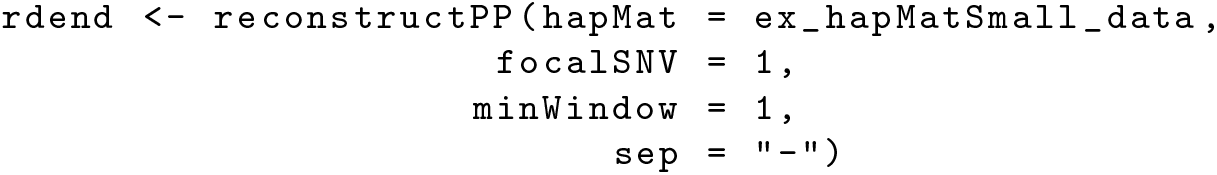

Figure 1 shows the reconstructed dendrogram, `rdend`, at the first SNV of `ex_hapMatSmall_data`. The structure of `rdend` is as follows:

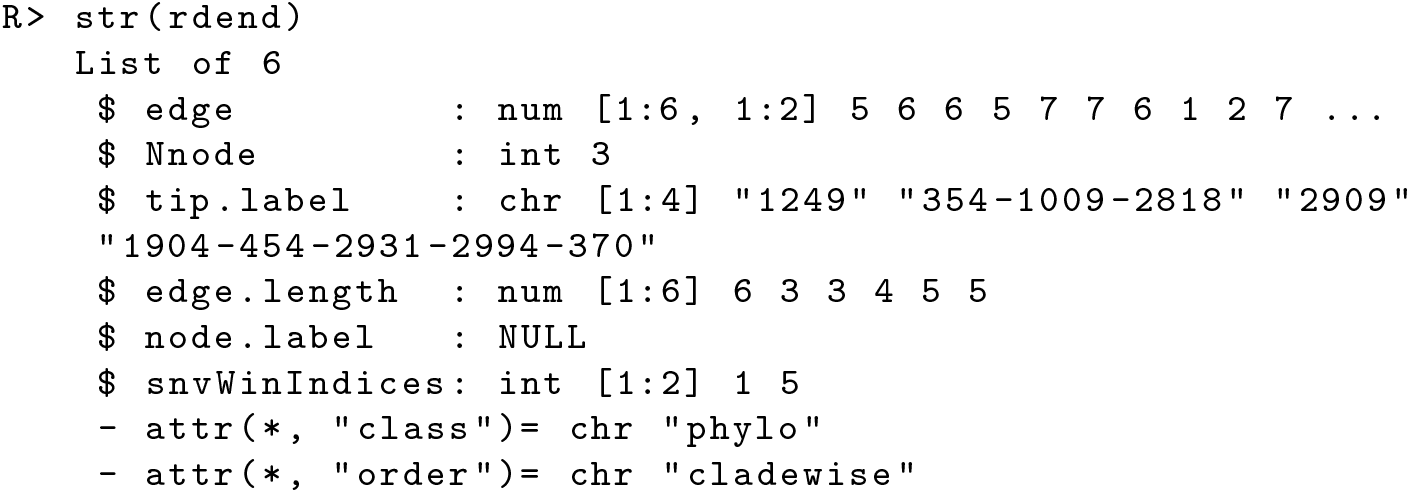

**Figure 1:**
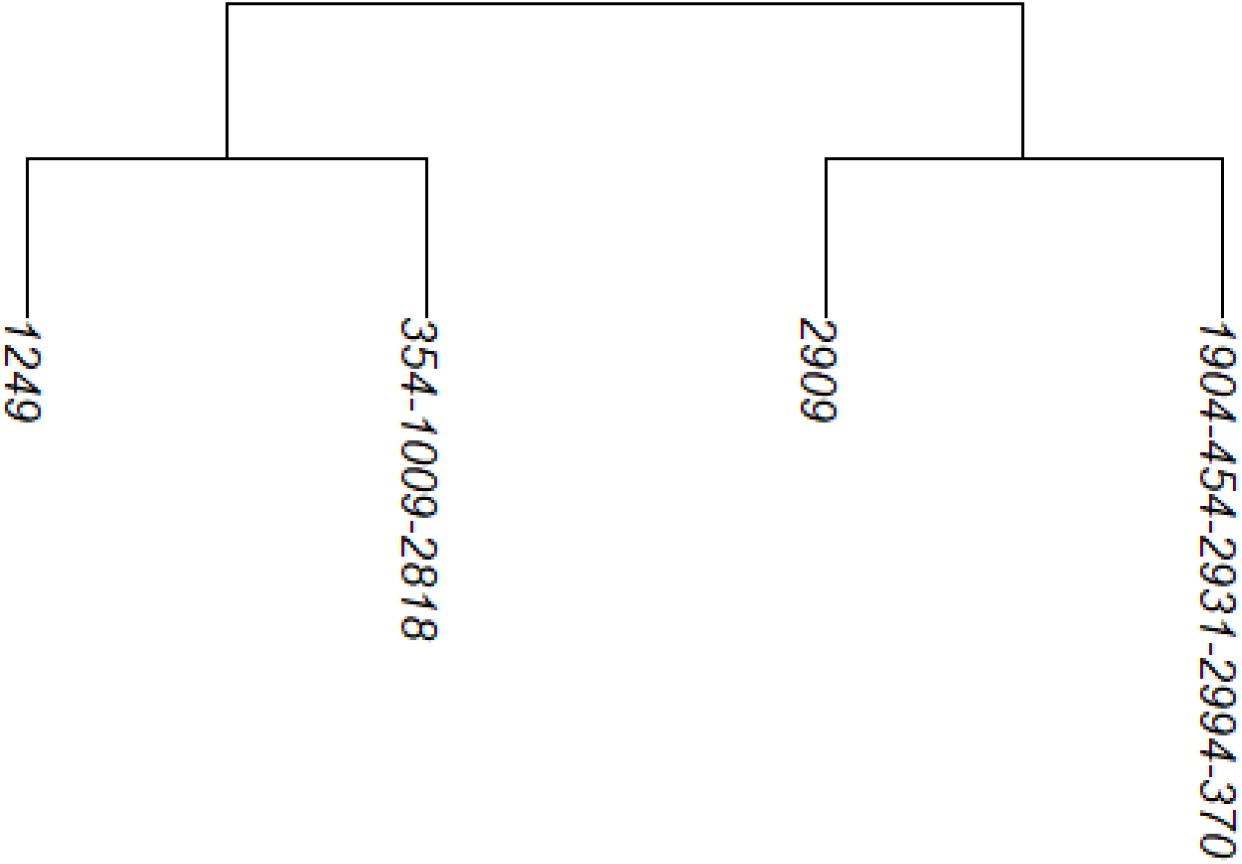
The reconstructed partition at the first SNV of `ex_hapMatSmall_data`.

The user can extract the positions of the lower and upper limits of the neighborhood of SNVs used to reconstruct `rdend` as follows:

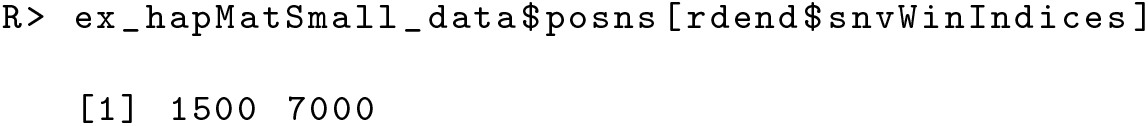

To see the sequences in the neighborhood of SNVs used for the reconstruction, the user can execute the following command:

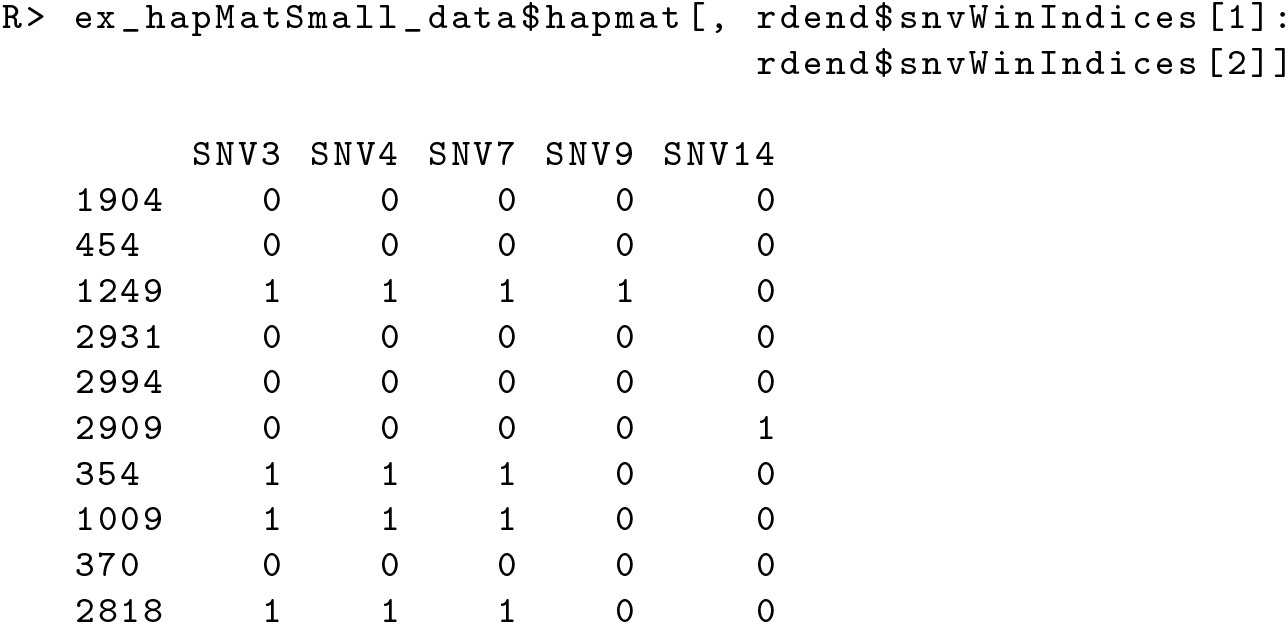

As can be seen in the above output, there are two groups of sequences that have the same ancestral and derived alleles at each SNV position: sequences 354, 1009 and 2818, and sequences 1904, 454, 2931, 2994 and 370. These two groups of sequences therefore cannot be distinguished in the reconstructed partition. In Figure 1, we can verify that two tips of the partition are comprised of these two groups of sequences.

With `reconstructPPregion()`, the user can reconstruct perfect phylogenies at each possible focal SNV in a `hapMat` data object. In the following example, we consider the 10 sequences with 20 SNVs in `ex_hapMatSmall_data`. We reconstruct perfect phylogenies across the 20 SNVs.

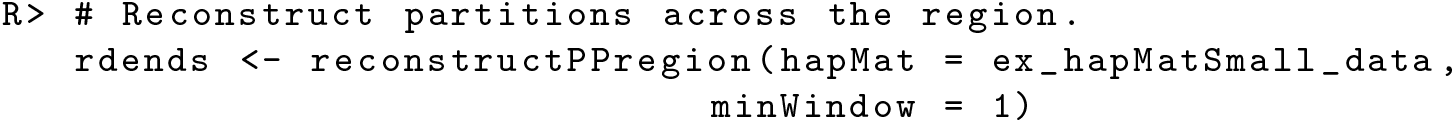

`rdends` is an `ape` multiphylo object. The reconstructed partition at the first focal SNV in `ex_hapMatSmall_data` is the first `phylo` object in `rdends`:

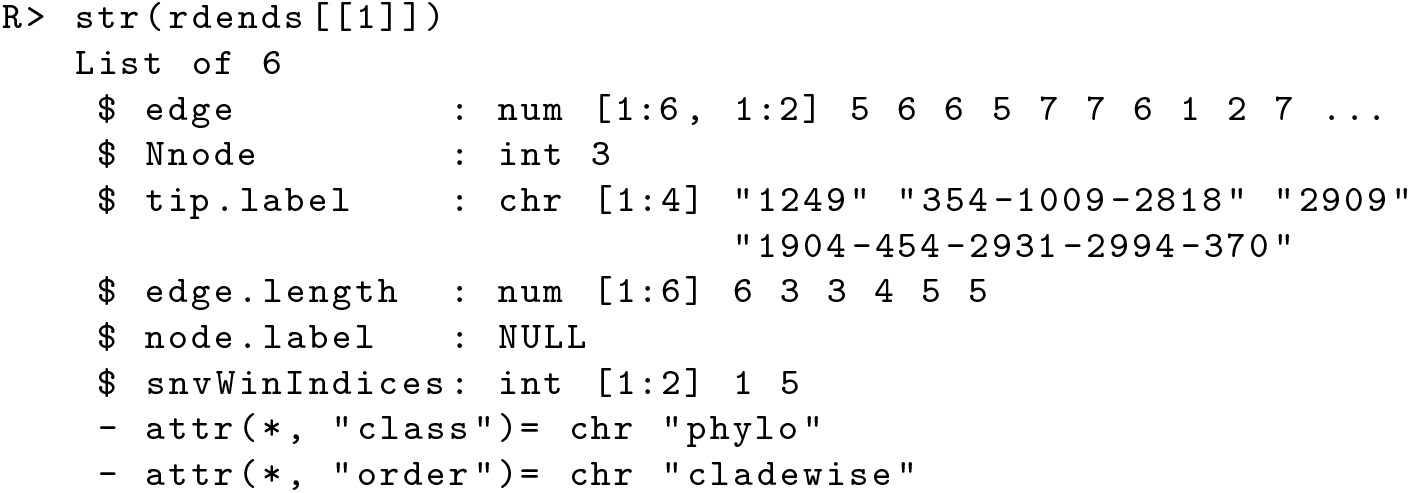

If a user wants to reconstruct perfect phylogenies within a user-provided subregion of a `hapMat` object, they may specify the lower and upper values of the subregion in base pairs as follows:

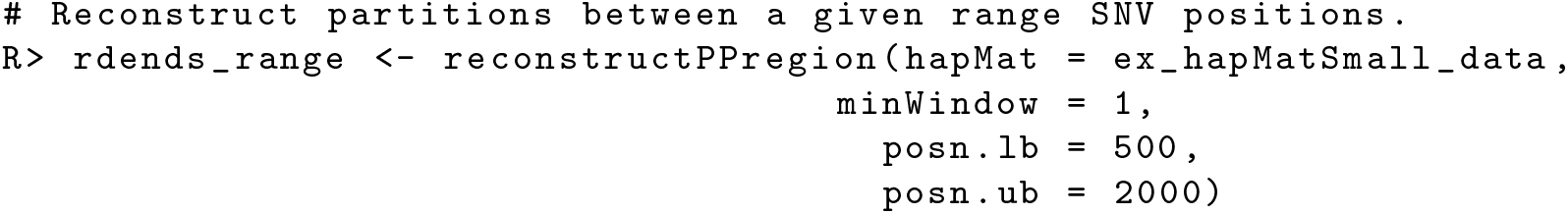

The function `testDendAssoRI()` uses the Rand Index to investigate the association between a comparator dendrogram or partition and multiple reconstructed dendrograms or partitions across a genomic region. Detailed descriptions of the function arguments and output of `testDendAssoRI()` are provided in the Additional file 1, along with a worked example.

Figure 2 shows the association profile between a comparator true dendrogram, `tdend`, at position 975 kbp, and a list of reconstructed dendrograms across the genomic region of `ex_hapMat_data`. In the two panels of the figure, the Rand indices are based on six and 24 clusters. Since we use simulated data, we know the true dendrogram at position 975 kbp. In Figure 2, using the Rand index, we investigate how the true dendrogram at position 975 kbp associates with the reconstructed dendrograms across the genomic region. As can be seen, the highest point for six clusters lies at position 975 kbp, and for 24 clusters is very close to position 975 kbp. According to the omnibus *p*-value, returned by `testDendAssoRI()`, the association across the genomic region is significant (*P* ≈ 0.001) for both six and 24 clusters.

**Figure 2:**
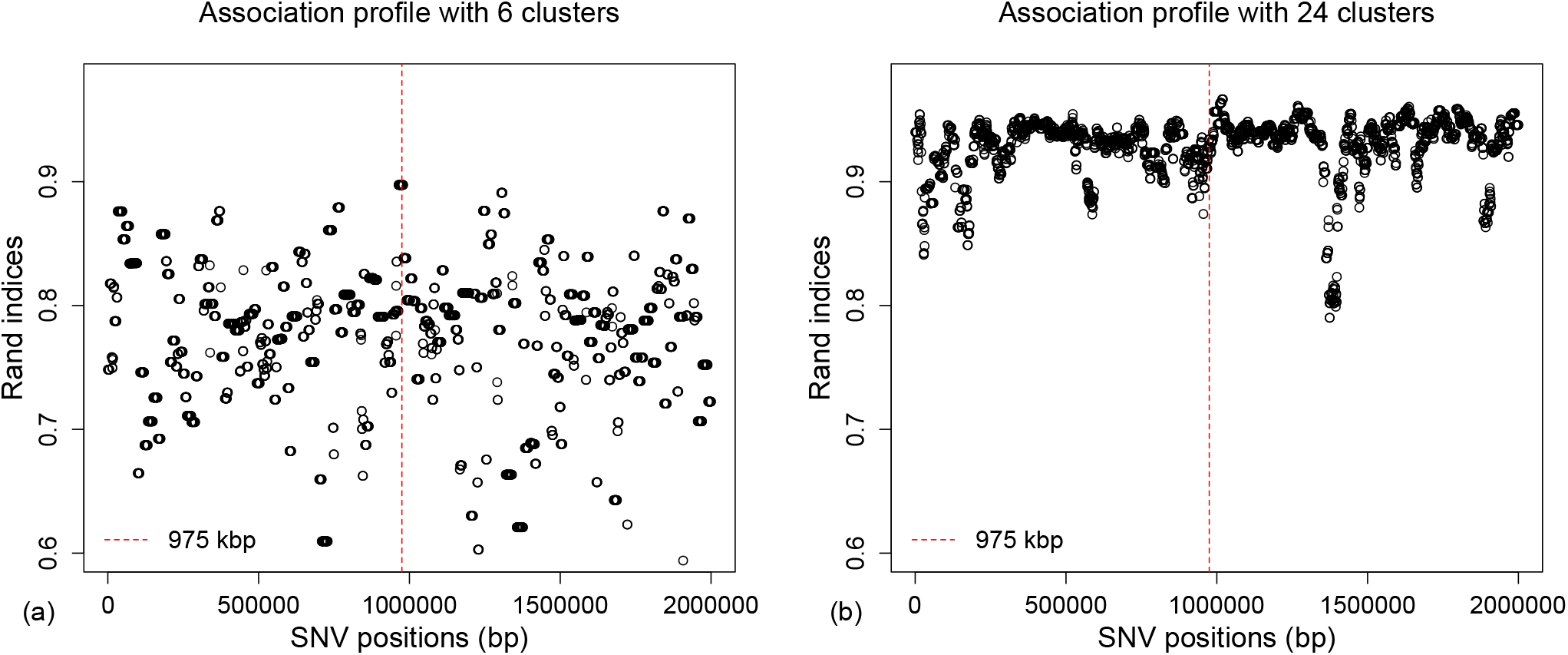
Rand indices associating a comparator true dendrogram at position 975 kbp and reconstructed dendrograms across the genomic region. **a)** Based on the six clusters. **b)** Based on 24 clusters. Red vertical dashed lines represent the position of the comparator dendrogram at 975 kbp.

The function `testAssoDist()` investigates the association between a comparator distance matrix and multiple reconstructed dendrograms across a genomic region. The association statistics available in the function are the dCor statistic, HHG statistic, Mantel statistic, and RV coefficient. The function has the following five key arguments:

1. `rdend`: An `ape` multiphylo object of reconstructed dendrograms at each focal SNV.
2. `cdmat`: A comparator matrix of pairwise distances (e.g. pairwise distances between sequences of a comparator dendrogram).
3. `method`: A character string specifying one of `“dCor”`, `“HHG”`, `“Mantel”` or `“RV”` for the dCor, HHG, Mantel or RV statistics, respectively.
4. `hapMat`: An object of class `hapMat` containing SNV sequences.
5. `nperm`: Number of permutations for the omnibus test of any association across the genomic region. The default is `nperm = 0`; i.e., association will not be tested.

To illustrate, we plot the dCor statistics summarizing the association between a comparator distance matrix, `cdmat`, and the reconstructed dendrograms across the genomic region of the example dataset `ex_hapMat_data`.

First, we compute the pairwise distances between sequences based on the comparator true dendrogram at SNV position 975 kbp. These pairwise distances are computed with the function `rdistMatrix()`, available in the package. The `rdistMatrix()` function uses the rankings of the nested partitions in the dendrogram to calculate rank-based distances between the sequences. However, users can provide any distance measures of interest for `cdmat`. We then plot the dCor statistic summarizing the association between the rank-based distance matrix for the reconstructed dendrograms at each SNV position and the comparator distance matrix at SNV position 975 kbp (Figure 3).

**Figure 3:**
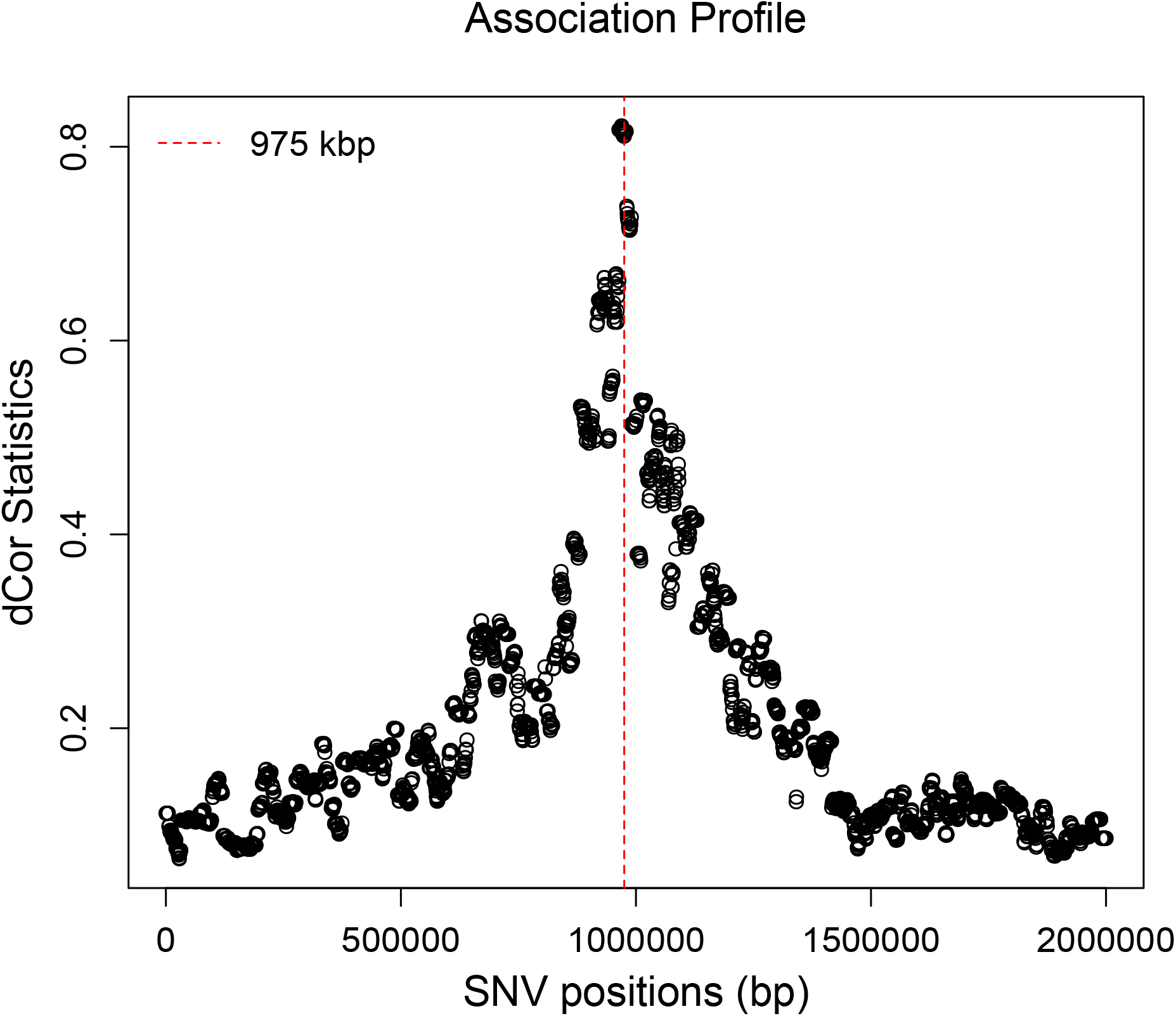
Associations between a comparator distance matrix from the true dendrogram at position 975 kbp and the reconstructed dendrograms across the genomic region. Red vertical dashed line represents the position of the comparator dendrogram at 975 kbp.

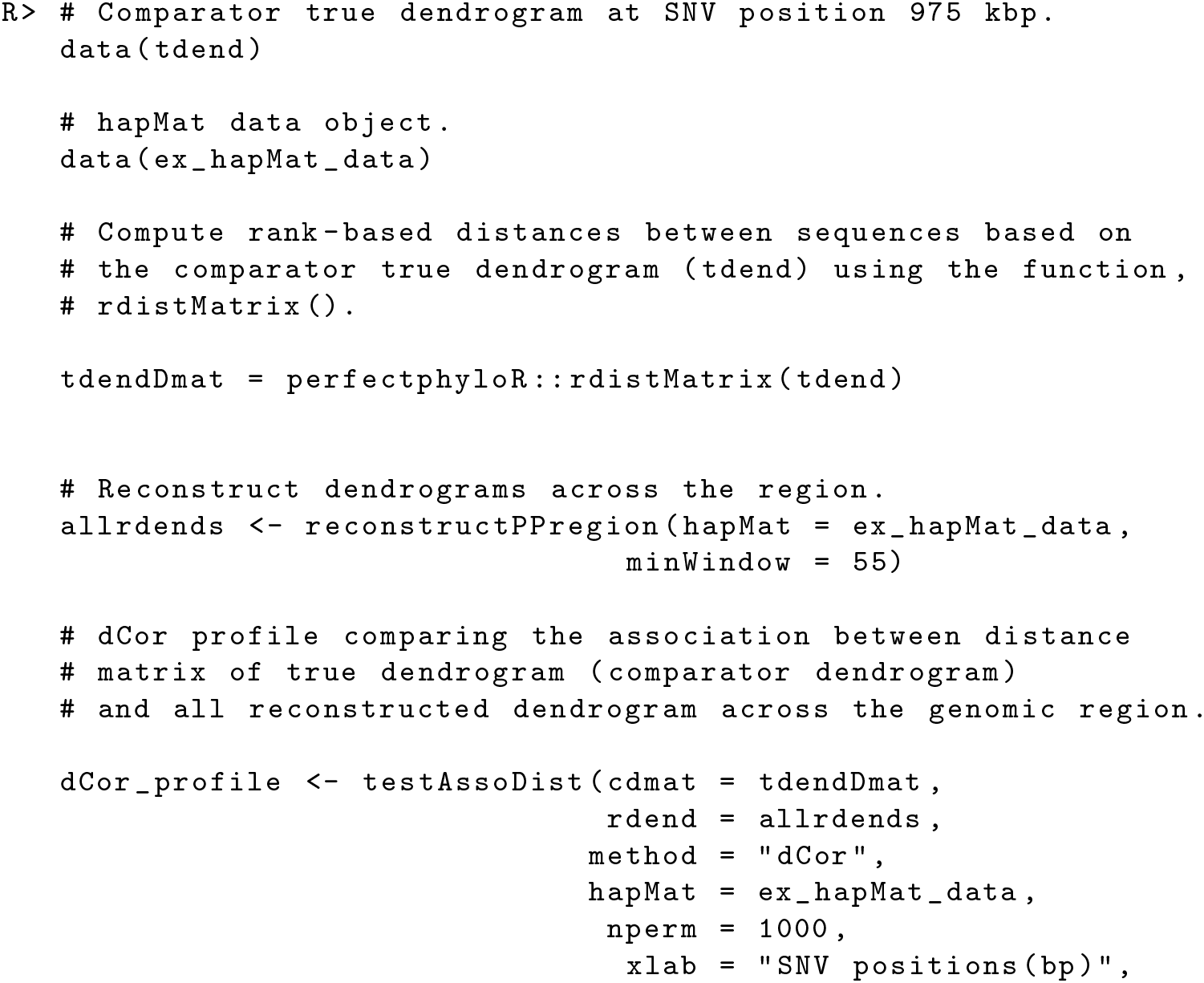

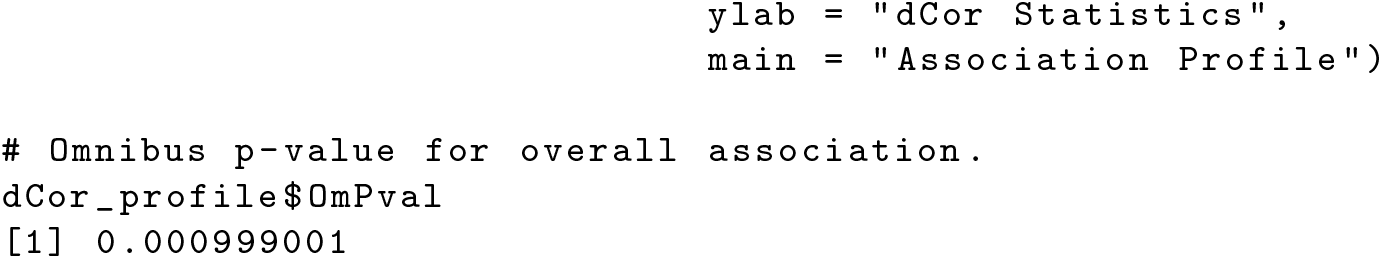

In Figure 3, we can clearly see the strongest association around the SNV position 975 kbp, and the association across the genomic region is significant (*P* ≈ 0.001), as expected. The association signal is much clearer than for the Rand index plotted in Figure 2 because dCor uses the full information from the pairwise distance matrices whereas the Rand index is based on a discrete number of clusters.

To illustrate another application of the function `testAssoDist()`, we perform the RV test of association between a phenotypic distance matrix as the `cdmat` argument and the reconstructed dendrograms across the genomic region of `ex_hapMat_data`. The phenotype data and distances are described in [19] and are contained in the data object phenoDist. Binary phenotype status was assigned based on causal SNVs from a causal subregion defined from 950 - 1050 kbp within the 2-Mbp genomic region.

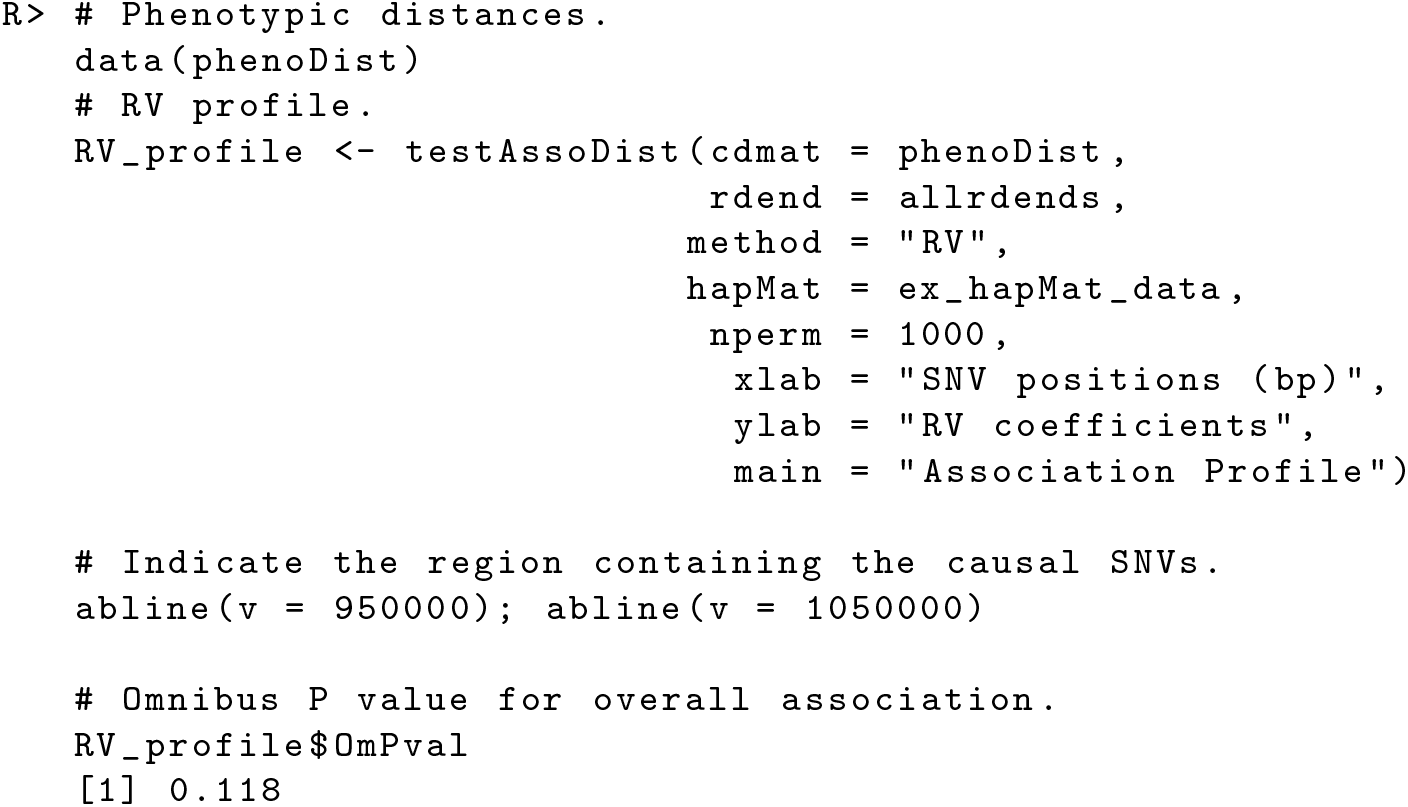

Figure 4 shows the resulting association profile between the phenotypic distances and the reconstructed dendrograms across the genomic region in `ex_hapMat_data`. The vertical lines indicate the causal subregion of 950 - 1050 kbp. The strongest association is close to the causal subregion. However, in this example, the association across the genomic region is not significant (*P* ≈ 0.1).

**Figure 4:**
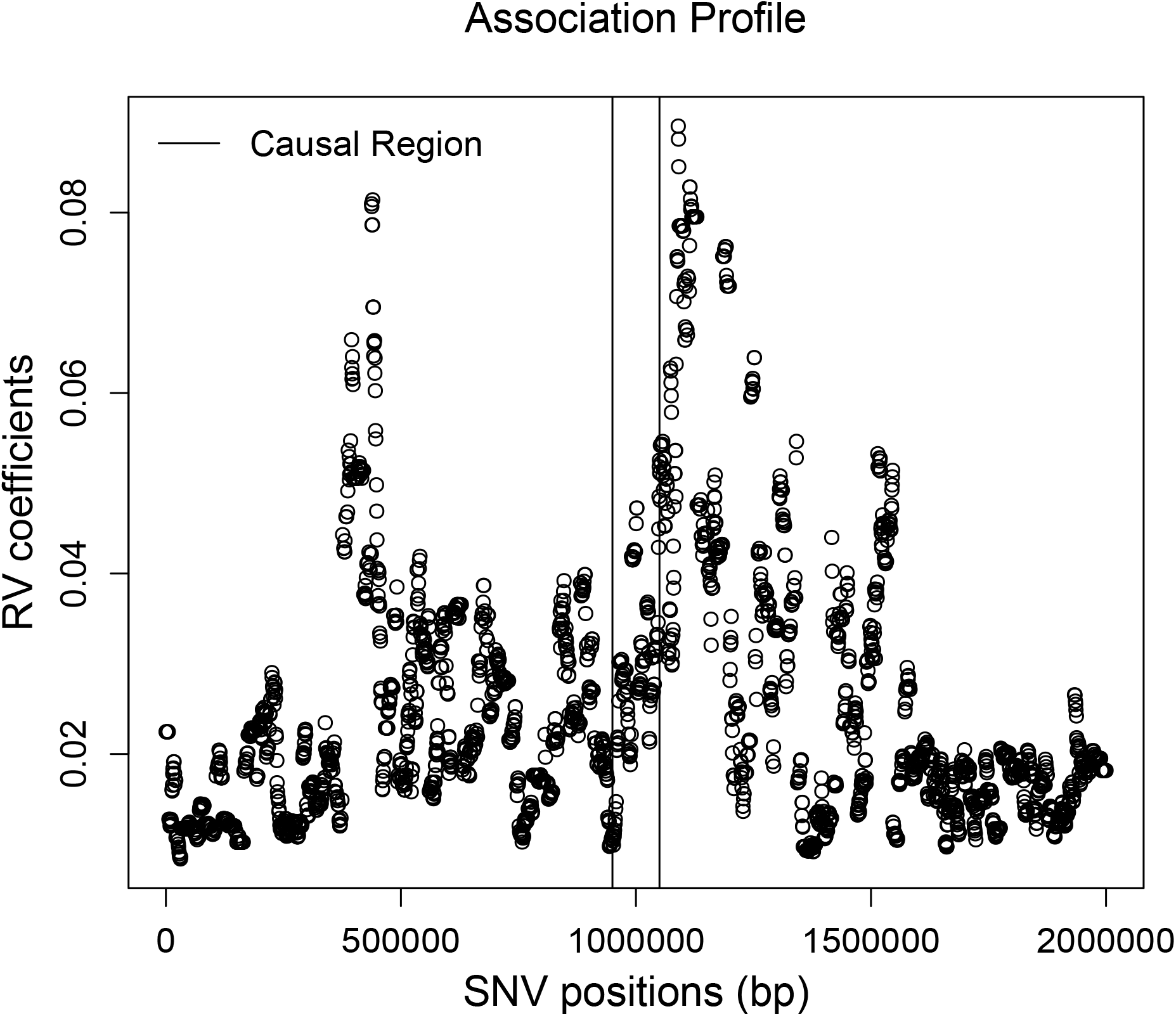
Associations between the phenotypic distance matrix and the reconstructed dendrograms across the genomic region. Black vertical lines indicate the limits of the genomic region containing trait-influencing SNVs.

## Timing

Table 1 shows the computation times of the package’s major functions. These computation times are for the 200 sequences comprised of 2747 SNVs in the example data `ex_hapMat_data` that is included in the package. Table 2 compares computation times of the function `reconstructPPregion()` for different numbers of sequences and numbers of SNVs. These times scale approximately linearly in the number of SNVs and quadratically in the number of sequences. Computation times are measured on an Intel E5-2683 v4 at 2.1 GHz with 20 GB of RAM.

**Table 1:** Computation times of the major functions of the package `perfectphyloR` for 200 sequences comprised of 2747 SNVs.

| Function | Computation Time (minutes) |  |
| --- | --- | --- |
|  | No permutation | 1000 permutations |
| `reconstructPP()` | Few seconds | NA |
| `reconstructPPregion()` | 12.00 | NA |
| `testDendAssoRI()` | 1.12 | 70.00 |
| `testAssoDist()` | 0.28 | 193.71 |

**Table 2:** `reconstructPPregion()` timing results (in minutes) for different number of sequences and SNVs.

| Number of SNVs | Number of sequences |  |  |  |
| --- | --- | --- | --- | --- |
|  | 200 | 300 | 400 | 500 |
| 3000 | 17.66 | 21.47 | 26.07 | 26.38 |
| 4000 | 23.38 | 28.26 | 33.41 | 33.80 |
| 5000 | 29.99 | 34.82 | 41.40 | 41.87 |
| 6000 | 36.58 | 43.41 | 49.86 | 50.67 |
| 7000 | 43.17 | 51.68 | 58.72 | 59.42 |
| 8000 | 49.25 | 59.11 | 67.89 | 68.65 |
| 9000 | 55.63 | 66.75 | 77.00 | 77.92 |
| 10000 | 61.56 | 75.53 | 86.11 | 86.99 |

## Discussion

We note that the computation time of `reconstructPPregion()` can vary a lot based on the size of the `hapMat` object (Table 2). Starting from the first SNV of the `hapMat` object, this function continues the reconstruction process until the last SNV. At each focal SNV, the function starts from ground level to construct a surrounding window of SNVs and rebuilds the partition, without utilizing the information from previously constructed partitions at nearby SNVs. As a result, many of the same computations may be done several times for similar focal SNVs. As noted by a reviewer, there may be ways to make `reconstructPPregion()` faster. For example, clustering similar successive SNVs before starting the reconstruction could lead to computational efficiencies and would be an avenue for future work.

Although we know of no software that is directly comparable to `perfectphyloR`, the PerfectPhy suite of tools is also set up to return sequence partitions. We therefore explored the use of PerfectPhy in a moving-window approach similar to that of `perfectphyloR`. Briefly, for each placement of the moving window, the following two steps were repeated: (i) filter out incompatible SNVs in the window and (ii) reconstruct the perfect phylogeny using the remaining compatible SNVs. We applied this approach to the 200 sequences in the example dataset, `ex_hapMat_data`, using the default minimum-window size of 55 for 2747 SNVs. For the first few window placements, we compared the computational time of steps (i) and (ii) in the PerfectPhybased approach to that of `reconstructPP()` in `perfectphyloR`. For the PerfectPhy approach, the filtering step is the bottleneck, with computation times in excess of 600 minutes. By contrast, `reconstructPP()` took no more than 0.18 seconds.

## Conclusion

The R package `perfectphyloR` provides functions to reconstruct a perfect phylogeny at a user-given focal SNV and perfect phylogenies across a genomic region of interest. The package also computes, tests and displays association measures based on the reconstructed partitions in a genomic region. The reconstructed partitions are useful to researchers seeking insight into the ancestral structure of DNA sequences. For example, associating the reconstructed partitions with a trait can help to localise trait-influencing variants in association studies. `perfectphyloR` can be freely downloaded from the Comprehensive R Archive Network (CRAN) or from https://github.com/cbhagya/perfectphyloR/.

## Availability and requirements

Project name: perfectphyloR

Project home page: https://CRAN.R-project.org/package=perfectphyloR

Operating system(s): Windows, Linux, OS X

Programming language: R

Other requirements: R 3.4.0 or newer

License: GPL-2, GPL-3

Any restrictions to use by non-academics: none

The package `perfectphyloR` can be installed from CRAN using `install.packages(“perfectphyloR”)`. The local zip file can be installed using R Studio by selecting the install package(s) from local zip files.

## List of abbreviations

DNA: Deoxyribonucleic acid
BLOSSOC: BLOck aSSOCiation
SNV: Single Nucleotide Variant
dCor: Distance Correlation
RI: Rand Index
HHG: Heller-Heller-Gorfin
RV: R-Vector, a vector version of standard *r* correlation
GHz: Giga Hertz
GB: Gigabyte
RAM: Random Access Memory
CRAN: Comprehensive R Archive Network

## Acknowledgements

We thank the anonymous reviewers for constructive comments that improved and clarified the manuscript. We also thank Christina Nieuwoudt and Kelly Burkett for helpful discussions and comments, and the Department of Statistics and Actuarial Science at Simon Fraser University for its generous support. This research was funded in part by a Discovery Grant from the Natural Sciences and Engineering Research Council of Canada.

## Appendix testDendAssoRI(): Arguments and example call

The function `testDendAssoRI()` has five key arguments:

1. `rdend`: An `ape` multiphylo object of reconstructed dendrograms at each focal SNV.
2. `cdend`: An `ape` phylo object of the comparator dendrogram.
3. `hapMat`: An object of class `hapMat` containing SNV sequences.
4. `k`: An integer that specifies the number of clusters that the dendrogram should be cut into. The default is `k` = 2. Clusters are defined by starting from the root of the dendrogram, moving towards the tips and cutting across when the appropriate number of clusters is reached.
5. `nperm`: Number of permutations for the test of any association across the genomic region. The default is `nperm` = 0; i.e., association will not be tested.

To illustrate, we use the example dataset `ex_hapMat_data` with 200 sequences and 2747 SNVs. We plot the Rand index values summarizing the association between the comparator dendrogram at SNV position 975 kilobase pairs and the reconstructed dendrogram at each SNV position across the 2 Mbp genomic region (Figure 2a).

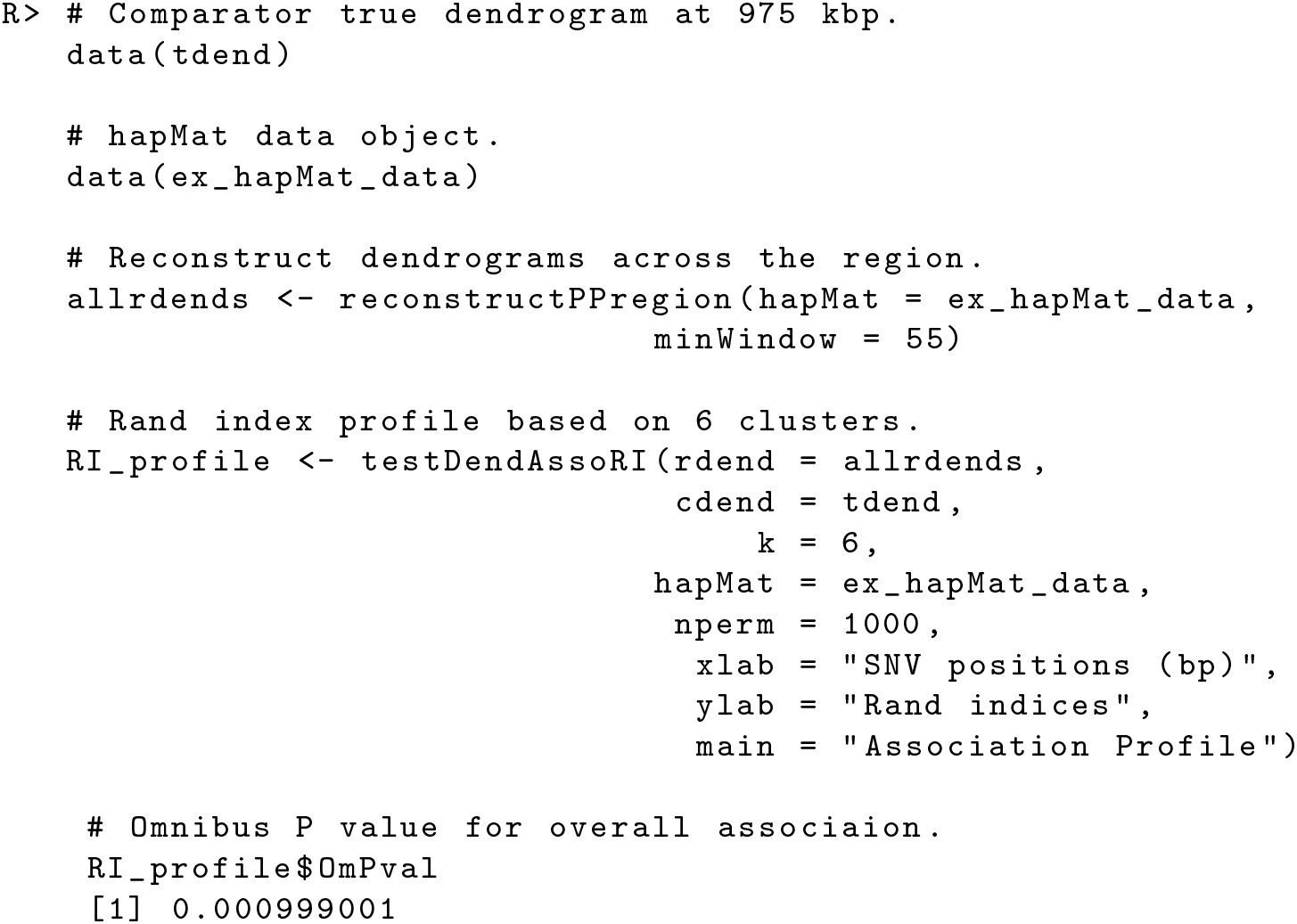

## Notes

https://github.com/cbhagya/perfectphyloR

